# Guiding clustering and annotation in single-cell RNA sequencing using the average overlap metric

**DOI:** 10.1101/2025.05.06.652497

**Authors:** Christopher Thai, Amartya Singh, Daniel Herranz, Hossein Khiabanian

## Abstract

Defining cell types using unsupervised clustering algorithms based on transcriptional similarity is a powerful application of single-cell RNA sequencing. A single clustering resolution may not yield clusters that represent both broad, well-defined populations and smaller subpopulations simultaneously. Therefore, when cell identities are not known prior to sequencing, robust comparison and annotation of inferred *de novo* clusters remains a challenge. In this work, we define the distance between single-cell clusters by proposing the use of the average overlap metric to compare ranked lists of differentially expressed genes in a top-weighted manner. We first benchmark our approach in a truth-known dataset comprised of highly similar yet distinct T-cell populations and show that evaluating clusters with average overlap results in a consistent, precise, and biologically meaningful recapitulation of true cell identities. We then apply our approach to data of unsorted mouse thymocytes and characterize stages of T-cell development in the thymus, including minor populations of double-negative (CD4-CD8-) T-cells that are notoriously difficult to confidently detect in unsorted single-cell data. We demonstrate that measuring cluster similarity with average overlap of marker gene rankings enables robust, reproducible characterization of single cells and clarifies biological interpretation of their underlying identities in highly homogeneous populations.

## INTRODUCTION

Understanding complex biological processes, such as hematopoiesis or tumorigenesis, requires an accurate identification of the types of cells present in a tissue sample and a mapping of their interactions with one another. Bulk RNA sequencing methods measure average gene expression across all cells^1^ and cannot measure cell-to-cell gene expression variation that may arise due to functional differentiation, such as those during T-cell development, or time-dependent processes that occur across tumor clonal evolution. Meanwhile, single-cell RNA sequencing (scRNA-seq) technologies have been developed to address these challenges, leading to an improved understanding of molecular processes in complex diseases^2^.

Defining cell types using unsupervised clustering algorithms based on transcriptional similarity is a powerful application of scRNA-seq. There are several steps involved in the computational analysis of scRNA-seq data^2–4^, including initial quality control, normalization, clustering, and identifying differentially expressed genes, which are considered as marker genes^5^ for each inferred cluster. One can then analyze these marker genes and consult the literature and reference databases to assign biological identities to each cluster. These steps are currently implemented as a complete workflow in commonly used analytical packages like Seurat^6, 7^ and Scanpy^8^.

Still, challenges remain in clustering of scRNA-seq data stemming from the nature of measurements from single cells^9^. Cell identities are not known before sequencing without a prior sorting process, and noise in measuring gene expression, especially for those expressed in small subpopulations, can confound cell clusters inferred by unsupervised community detection algorithms. Additionally, parameters that determine the number and membership of clusters, such as the resolution in Leiden or Louvain methods^10, 11^, are applied globally across an entire dataset, despite the possible presence of distinct subpopulations with varying sizes. To avert over- or under-clustering and subsequent mischaracterization of novel cell populations and their marker genes, one can directly annotate cells using their expression of pre-defined sets of marker genes^12–15^. However, direct annotation is constrained to identities present in reference datasets and may miss novel biological populations uncharacterized in the literature. Meanwhile, solutions for optimizing clustering parameters have been proposed^16–20^, but consistent biological interpretation of inferred clusters remains an unresolved necessary step.

Inferring similar cell state or identity for clusters that share key marker genes is analogous to a subjective call on two lists’ similarity based on the presence of a few key members. Conversely, the mere presence of the same genes in two or more clusters’ marker gene sets may not by itself indicate the same cell state or identity. Differences in the magnitude of differential expression in each cluster, and by extension, differences in the rankings of marker genes, may indicate a relevant differentiation trajectory or novel subpopulation. Yet, methods are lacking for the quantification of marker gene list similarity between single-cell clusters to annotate cell identity.

Metrics that compare two lists based on the ranking of their elements address this challenge by providing a measure to compare single-cell marker gene lists from cluster-based differential expression analyses. While rank-based methods for scRNA-seq analyses have been developed^21–23^, none specifically address cluster similarity based on differential expression, especially when clusters are computationally determined in an unsupervised manner through community detection algorithms like Leiden or Louvain. To date, no quantification of the similarity of differentially expressed genes and their rankings has been employed in downstream scRNA-seq analysis. Here, we provide a definition for similarity between single-cell clusters based on a rank-based metric called average overlap^24^ and show that it can be used to calculate the significance of related cell identities in a truth-known dataset with consistency, precision, and meaningful biological interpretation. We then demonstrate the utility of average overlap for guiding the clustering and annotation of cells in the thymus, revealing individual stages of thymocyte development.

## RESULTS

### Average overlap metric for cluster marker gene comparison

To compare single-cell clusters derived from unsupervised community detection algorithms, we first defined each cluster’s marker gene set by its most differentially expressed genes relative to the rest of the cells. We then ranked the marker gene set based on the significance of differential expression and hypothesized that clusters that share marker genes with similar rankings are more likely to have similar cell identity and/or state. To test this hypothesis, we used a rank-based metric called average overlap (AO)^24^, which is a top-weighted measure of ranked list similarity. In the context of marker gene sets, AO weighs differences in the rankings of the most differentially expressed genes higher than for the genes lower in the set. AO is defined by the mean of the overlaps between two ranked lists calculated at a range of depths into the lists (**Fig. 1a**). AO distances range from 0 to 1, from completely dissimilar to completely identical lists.

**FIGURE 1.**
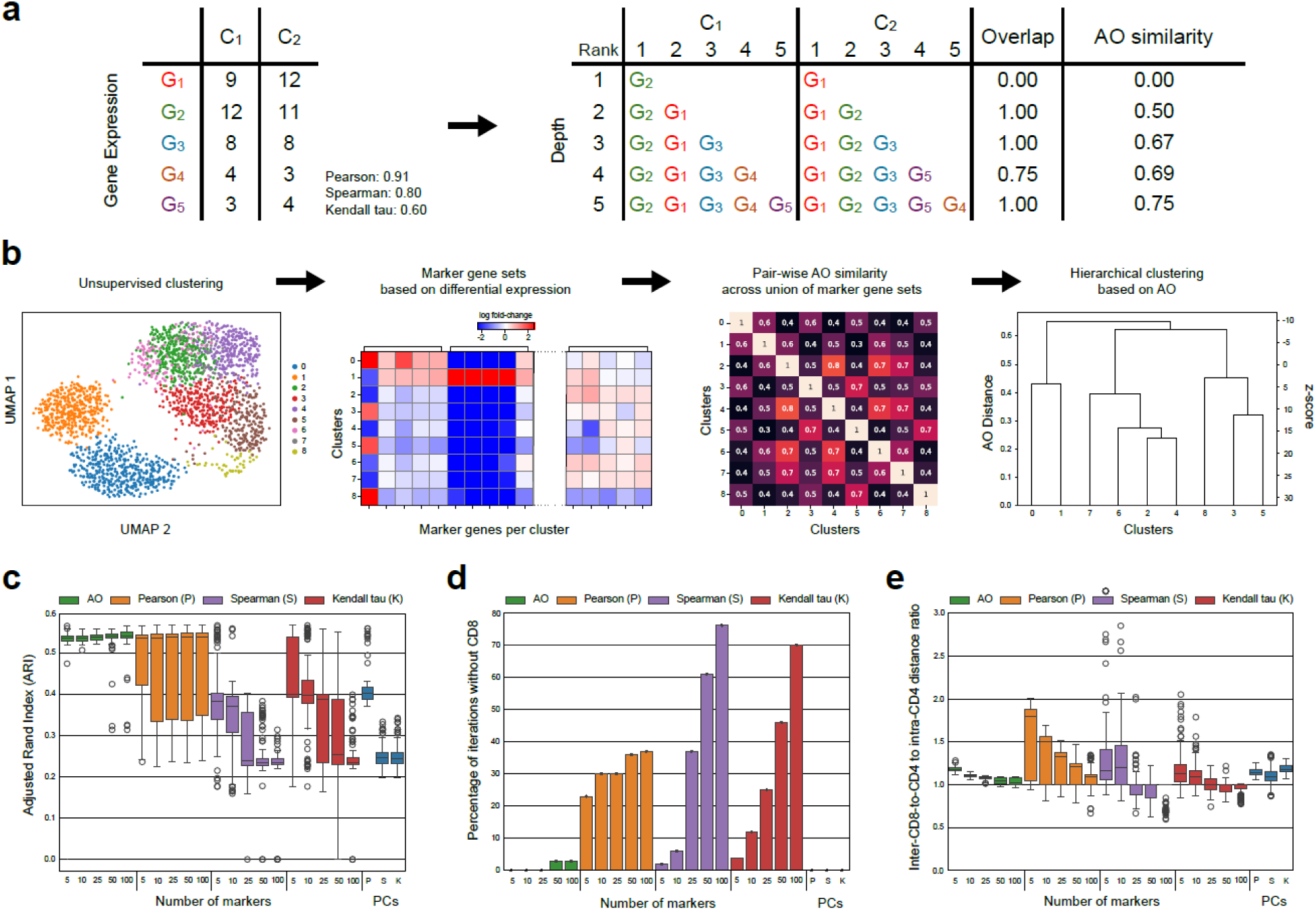
Workflow and benchmarking of average overlap for quantitative comparison of inferred single-cell clusters in scRNA-seq data. **a**, Calculating the average overlap distance between two ranked lists and comparison with correlation-based metrics. Differentially expressed genes (G1 to G5) are ranked in clusters C1 and C2 and the overlap between the ranked lists is calculated at each depth. Average overlap of the entire lists is the mean of all overlaps. **b**, A schematic demonstrating the workflow for hierarchical clustering using average overlap, following community detection, marker gene identification, and pair-wise distance calculation. **c**, Adjusted Rand Index (ARI) performance measures based on agreement with ground truth labels in five derived cell populations, across different metrics and number of marker genes used, using the T-cells in the *Zhengmix8eq* dataset. Adjusted mutual information (AMI) and the Fowlkes-Mallows score (FMS) are shown in **Supplementary Fig. 2**. **d**, Percentage of benchmarking trials in which a final cluster enriched with ground-truth CD8 naïve cytotoxic population was identified. **e**, Ratio of CD8 to CD4 distances versus intra-CD4 distances. A ratio higher than 1 indicates higher degree of separation between CD8 cytotoxic and CD4 populations versus the separation among the CD4 populations.

When calculated over randomly shuffled lists, AO closely follows a normal distribution (**Supplementary Fig. 1**) which provides a statistical means for assigning significance to AO scores and calculating the likelihood that two clusters share the same cell identity.

### Benchmarking the average overlap metric in truth-known data

To evaluate AO for quantifying differences between highly similar cell types, we used T-cell populations from a dataset with truth-known, biological cell identities, including equal numbers of purified T-helper, memory, naive, and regulatory CD4 T-cells as well as cytotoxic CD8 T-cells^20, 25^. We refer to this dataset as *Zhengmix8eq*.

We used the Piccolo workflow^26^ to process and normalize the raw counts and the Leiden method^11^ (resolution = 2.0) to produce single-cell clusters. After determining each cluster’s top-k marker genes (k = 5, 10, 25, 50, and 100), we calculated pairwise AO distances between the Leiden clusters based on their rankings of the union of all marker gene sets. To obtain five cell populations that corresponded to the five present T-cell populations in *Zhengmix8eq*, we performed hierarchical clustering of the initial Leiden clusters using the pairwise AO distances, and then iteratively merged the clusters with the highest AO similarity (**Fig. 1b**). We also labelled each final cluster based on the enrichment of ground truth labels and quantified the agreement between the five derived cell populations and known T-cell identities using the adjusted Rand index (ARI) **(Fig. 1c)**, adjusted mutual information (AMI), and the Fowlkes-Mallows score (FMS) (**Supplementary Fig. 2**). We repeated this analysis 100 times with a new random seed for running the Leiden method. To benchmark AO against other metrics utilized for assessing similarities between single-cell clusters, we performed the same hierarchical clustering with pairwise Pearson, Spearman, and Kendall Tau correlations, using the marker genes’ normalized expression counts. We also benchmarked the use of correlation metrics applied to the first 50 principal components, which summarize expression across all genes.

Five cell populations obtained from merging the Leiden clusters based on AO similarity showed the highest correspondence to true T-cell populations present in the dataset compared to all other metrics (**Fig. 1c**). Furthermore, AO reproducibly separated a CD8 cytotoxic T-cell population from the other CD4 T-cell populations. AO was able to distinguish a CD8-enriched population in 100% of iterations with 5, 10, and 25 marker genes and in 97% of iterations with 50 and 100 maker genes. In contrast, CD8-enriched populations were not detected when we merged the Leiden clusters with Pearson, Spearman, or Kendall Tau correlation of marker genes’ expression counts in 63-77%, 24-94%, and 30-96% of iterations, respectively (**Fig. 1d**).

Secondly, AO similarity consistently measured larger distances between the CD8 and CD4 populations than the pair-wise distances among the CD4 sub-populations. A ratio of the inter-CD8-to-CD4 distances versus the intra-CD4 distances that is greater than one indicates larger separation of the CD8 group and the CD4 groups. When using AO, the distance ratios were consistently greater than one across all marker gene numbers of 5 (range: 1.12-1.28, standard deviation (s.d.): 0.029), 10 (range: 1.06–1.15, s.d.: 0.018), 25 (range: 1.015–1.10, s.d.: 0.022), 50 (range: 0.97–1.10, s.d.: 0.034), and 100 (range: 0.96–1.09, s.d.: 0.039). In comparison, correlation-based methods showed substantially poorer performance when applied to marker gene expression counts. Pearson, Spearman, and Kendall-Tau distances distinguished CD8 populations from CD4 sub-populations in the same manner of distance ratios greater than one in only 75%, 81%, and 80% of iterations respectively at their best performance, which was achieved with 5 marker genes per cluster (Pearson range: 0.94–2.0, s.d.: 0.38; Spearman range: 0.89–5.09, s.d.: 0.65; Kendall Tau range: 0.84–2.05, s.d.: 0.24) (**Fig. 1e)**.

Measuring single-cell cluster similarities using a smaller number of marker genes per cluster resulted in better performance for AO and other tested rank-based metrics. Curiously, although Spearman and Kendall-Tau correlations were able to recover CD8 populations distinct from CD4 sub-populations in 100% of iterations when they were applied to 50 principal components (PC) (**Fig. 1e**), merging Leiden clusters using PC-based distances resulted in the weakest correspondence to true T-cell labels present in the dataset (**Fig. 1c**).

Put together, AO robustly and reproducibly measured similarities between single-cell clusters based on the ranking of marker genes and significantly outperformed correlation-based metrics in identifying and characterizing T-cell populations present in *Zhengmix8eq*.

### Thymic T-cell development at single-cell resolution

Specific stages of T-cell development in the thymus have been characterized with single-cell transcriptomic studies, starting from double negative (DN) populations DN1-DN4, advancing to immature single positive (ISP) and double positive (DP), and eventually mature CD4 and CD8 T-cells. While the T-cell development trajectory in the thymus has been partially resolved in normal and diseased states, characterizing double negative cell subpopulations using single-cell approaches has been especially challenging, in particular when measuring total thymocytes without prior cell sorting, leading to grouping of all DN subpopulations together without specifically differentiating DN1-DN4 states^27–30^. To address this challenge and to demonstrate the utility of the AO metric in elucidating the relationships between thymic cell populations, we used our methodology to analyze single-cell transcriptomic data previously published from mouse thymocytes^28^. We processed the data using the Piccolo workflow and used the Leiden method for cell clustering (resolution = 1.0), obtaining 19 single-cell clusters (**Fig. 2a**). Marker gene sets for each single-cell cluster were defined by performing differential gene expression analysis relative to all other cells. We then combined each cluster’s marker gene set, composed of its top 25 genes ranked by significance, and performed hierarchical clustering using pairwise AO distances between each cluster’s rankings of the marker genes, testing the hypothesis that T-cell clusters with high AO correspond to cell populations with similar identity and state (**Fig. 2b,c**). To guide annotating T-cell identities for unsupervised clusters, we utilized the MyGeneSet tool, available as a part of the Immunological Genome Project (ImmGen)^31^ and mapped the expression of each cluster’s top 50 marker genes across 12 sorted mouse thymic cell populations using composite z-scores (**Fig. 2d**). This analysis linked the expression profiles of the 19 *de novo* single-cell clusters to MPP4 (a subset of multipotent progenitors from the bone marrow), DN1, DN2a, DN2b, DN3, DN4, ISP, DP, mature CD4 and CD8, natural killer, and γδ T-cells in ImmGen cell populations^32^.

**FIGURE 2.**
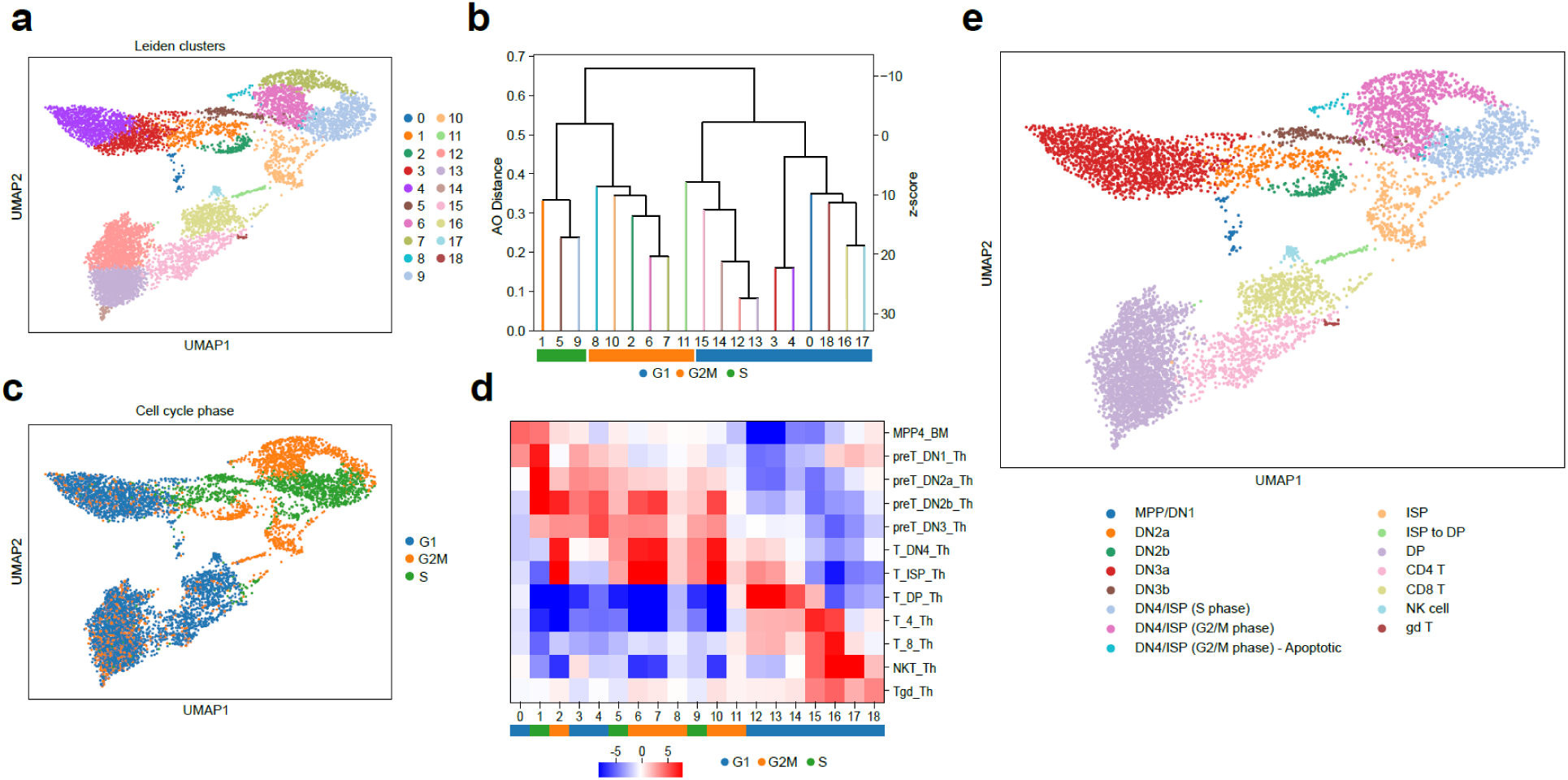
Utilizing average overlap to characterize stages of T-cell development in a mouse thymus. **a**, A UMAP plot of mouse thymocytes showing 19 inferred clusters using the Leiden algorithm. **b**, Cluster tree generated from hierarchical clustering of Leiden clusters with average overlap of marker gene rankings, based on the combined set of each cluster’s top 25 marker genes. Clusters largely group based on differential expression of genes related to the cell cycle. **c**, A UMAP plot of the mouse thymocytes, where each cell is annotated with its inferred cell cycle phase. **d**, A heatmap summarizing expression of Leiden cluster marker genes in sorted bulk populations of thymocytes from ImmGen. Each cluster is also colored with its inferred cell cycle phase. **e**, Final annotations of developing mouse thymocytes.

### The cell cycle characterizes thymic single-cell clusters

Marker genes in multiple single-cell clusters included those related to phases of the cell cycle. Hierarchical clustering based on AO of marker gene sets identified distinct groups of single-cell clusters; therefore, we hypothesized that similarity in the rankings of differentially expressed genes, as quantified by AO, would group the clusters based on cell cycle phases. We tested this hypothesis by scoring each cell based on the expression of known cell cycle genes and predicting their cell cycle phase^33^ (**Fig. 2b**). We found that clusters 1, 5, and 9 were enriched with cells in the S phase while clusters 2, 6, 7, 8, 10, and 11 were enriched with cells in the G2/M phase (**Fig. 2c**). The remaining clusters were enriched with cells in the G1 phase, split into two groups of clusters 0, 3, 4, 16, 17, and 18 plus clusters 12, 13, 14, and 15 (**Fig. 2c**). These marker genes associated with the cell cycle were also highly expressed in multiple ImmGen bulk populations, including the DN2b, DN4, and ISP populations (**Fig. 2d**). These results suggest that the groups that emerge from AO clustering primarily reflect cell cycle signatures known to play crucial roles in the identities of thymic cell populations, particularly pointing to stages of T-cell development involving rapid expansion and proliferation such as those that occur after T-lineage commitment and β-selection^34^.

### Annotation of thymic single-cell clusters during development

Using the annotations from MyGeneSet and Immgen sorted populations, we found that the earliest thymocyte progenitors (MPP4 and DN1 cells) mapped most strongly to cluster 0, which had marker genes that did not show significant expression in any other ImmGen thymic population. At the same time, cluster 0 did not show high similarity to any other cluster according to AO; thus, we annotated cluster 0 as a mix of multipotent progenitors (MPP) and DN1 cells.

Cluster 1 corresponded strongly to DN2a populations and cluster 2 was inferred as DN2b cells based on its marker genes’ expression specifically in the DN2a/b and earlier progenitor populations. Marker genes for cluster 2 were also highly expressed in proliferating DN4 and ISP populations, stemming from a 36% overlap with known cell cycle genes, including *Hmmr* and *Nusap1*.

Clusters 3 and 4 marker genes were highly expressed in DN3 populations and showed marked AO similarity. In addition, marker genes for cluster 5 showed relatively high expression in DN3 cells. For deeper characterization of these cells into DN3a and DN3b populations, we scored cells in clusters 3, 4 and 5 against genes upregulated in purified wildtype DN3a and 3b cells^35^. While cluster 4 highly correlated with DN3a, and cluster 3 modestly correlated with both phases, cluster 5, which is in the S phase of the cell cycle, showed similarity to DN3b cells, together indicating a transition of cells from DN3a to DN3b states (**Supplementary Fig. 3**). Accordingly, we merged clusters 3 and 4 together to form a DN3a population, and annotated cluster 5 as the DN3b population.

Clusters 6, 7, 8, and 9 mapped strongly to ImmGen DN4 and ISP populations. While cluster 9 was distinguished as DN4/ISP cells in the S phase of the cell cycle, clusters 6, 7, and 8 grouped together based on their pairwise AO and association with the G2/M phase of the cell cycle. Of note is cluster 8, which contained cells with low expression of a select set of genes that were not expressed in any other clusters (**Supplementary Fig. 4**). Relative to cells in clusters 6 and 7, cells in cluster 8 showed an upregulation of mt-Co1, mt-Co2, mt-Co3, and other mitochondrial genes. In fact, mitochondrial genes made up for on average 7.8% of this cluster’s expressed transcripts, indicating that a portion of cluster 8 would fall just under the 12% mitochondrial expression threshold we used for filtering low-quality cells at the start of analysis. Put together, we interpreted cluster 8 as representing apoptotic DN4/ISP cells (**Supplementary Fig. 5**).

Cluster 10 showed the highest correspondence to ImmGen ISPs, while cluster 11 was inferred as an intermediate population between the ISP and DP stages based on its marker genes’ expression in late development T-cells (**Supplementary Fig. 4**). Clusters 12, 13, and 14 were identified as parts of a larger DP population, as they each mapped strongly to ImmGen DP cells and showed the highest AO similarities across all clusters. The maturing stages of T-cell development were represented by cells in clusters 15 as single positive CD4 and cluster 16 as single positive CD8 T-cells. Finally, clusters 17 and 18 were assigned to smaller groups of natural killer cells and γδ T-cells, respectively.

Lastly, we explored characterizing the stages of T-cell development in our data using orthogonal methods. First, we asked whether pseudotime inference could recover cell identities in these developing T-cells and used diffusion pseudotime^36^ specifying cluster 0 as the starting point. The inferred trajectory matched our annotated stages of development and supported later developmental trajectory for cells in clusters 6 and 7 compared to cells in cluster 2; hence, partially distinguishing DN2b from DN4/ISP cells despite being in the same cell cycle phase **(Supplementary Fig. 6a**). This pattern, however, did not hold when cluster 2 was chosen as the analysis’s starting point **(Supplementary Fig. 6b)**. Cells in clusters 6 and 7 were assigned the same starting pseudotime value of 0 as the chosen cluster 2, indicating that pseudotime calculations can be confounded by strong cell cycle signatures. Second, we asked whether Pearson correlation applied to the expression counts of the marker genes could infer T-cell similarities. The resulting hierarchical clustering, however, did not consistently group the single-cell clusters based on phases of cell cycle or stages of T-cell development (**Supplementary Figure 7**), highlighting the strength of AO in inferring biologically meaningful relationships by using the rankings of differentially expressed genes as opposed to their counts, especially in the context of highly similar cells.

Overall, our extended analyses of truth-known hematopoietic populations as well as unsorted mouse thymocytes showed that AO could accurately guide the clustering and annotation of highly homogenous cell populations and help characterize the trajectory of T-cell development in thymus, including the detection of elusive double-negative (CD4-CD8-) cells. We have implemented the AO metric in Python and have published the ‘sc_average_overlap’ Python package, which is available at https://github.com/chrisvthai/sc_average_overlap, designed to work seamlessly within the Scanpy framework.

## DISCUSSION

In this work, we propose using average overlap, a top-weighted metric that quantifies the similarity of ranked lists, to compare clusters in scRNA-seq analysis. Hierarchical clustering using rank- and correlation-based metrics to compare the transcriptomic profiles of inferred clusters in scRNA-seq analysis is not new. Current implementations, including those in Seurat and Scanpy, utilize distances based on gene expression counts or principal components. In contrast, AO measures cluster similarity by relying on marker gene rankings derived from differential gene expression analysis.

We compared AO to correlation-based metrics using Leiden clusters derived from a biological, ground-truth dataset, containing five transcriptionally similar T-cell populations^20, 25^. We showed that merging unsupervised clusters into five groups with AO based on gene rankings resulted in cell labels that corresponded to ground truth populations more accurately than other tested methods. AO is also far more consistent in its performance, and over many Leiden clustering iterations, produced merged populations that were biologically meaningful, by differentiating CD8 cytotoxic populations from the CD4 clusters. Expression counts across highly similar populations are highly correlated irrespective of the feature selection procedure and the amount of cluster marker genes used. This high transcriptional correlation often results in inconsistencies that make distinguishing subtle differences between T-cell subpopulations and inferring cell clusters that map back to individual subpopulations a challenge. Restricting analysis to just gene rankings based on differential expression in our implementation of AO allows robustly measuring subtle transcriptional differences between single-cell clusters.

Moreover, AO performed well compared to other distance metrics due to its top-weighted property. While performance for all metrics diminished as more marker genes were included, AO-based hierarchical clustering was minimally affected. Reincorporating information about gene expression variation by way of utilizing principal coordinates recovered the performance of correlation-based metrics to an extent. We interpret this observation as evidence for the effect commonly known as the curse of dimensionality^37^ where the differences in distances between clusters become increasingly similar as a higher number of marker genes are added. In contrast, differences at the top of rankings as measured by AO, correspond to the most differentially expressed genes, and reflect cellular differences more precisely than the genes at the bottom of the rankings. As such, AO is not as sensitive to the curse of dimensionality as the other benchmarked metrics.

AO can quantify subtle differences in otherwise highly transcriptionally correlated cell populations. In our thymus data, AO-based hierarchical clustering revealed cell cycle genes as the strongest drivers of marker gene similarity. However, showing high correlation with ImmGen reference expression profiles was not sufficient to characterize cell cycle signatures in single-cell clusters that corresponded to DN2b, DN4, and ISP cells at once. By measuring differential gene rankings, AO inferred cell cycle phases with great specificity and guided identification of stages of T-cell development involving increased expansion and proliferation. Additionally, AO helped characterize a small subset of cells in the G2/M phase as a population of apoptotic DN4/ISP. Cells in this cluster were retained simply due to the choice of mitochondrial expression threshold used to account for low quality cells, yet showed relatively high correspondence to ImmGen DN4 and ISP populations, supporting their biological relevance. Thymocytes may undergo positive selection based on their ability to bind to MHC ligands; thymocytes unable to bind MHC undergo death by neglect^38^. The combination of relatively high mitochondrial gene expression which typically indicates dying cells, high correspondence to bulk DN4 and ISP populations in ImmGen, and the detection and quantification of unique marker genes via the Leiden clustering algorithm and the AO metric supports this interpretation. Altogether, unique subpopulations of thymocytes characterized with the aid of AO may be missed when relying solely on reference transcriptomic profiles.

The analysis and comparison of cluster marker genes are necessary steps in uncovering novel cell populations. As we have demonstrated with our analysis of thymus data, one clustering resolution may not yield clusters that simultaneously represent broad well-defined populations as well as smaller novel subpopulations. Moreover, while some clusters may separate due to existing biological variations, others may arise due to noisy clustering. AO-measured distances based on the ranking of differentially expressed genes provide a quantitative method for evaluating transcriptional similarity independent of clustering algorithms and parameters used. In the context of differentiating cells, alternative methods such as pseudotime inference, which calculates the relative position of a cell across gene expression gradients, can theoretically aid cluster annotation^36, 39, 40^. However, pseudotime algorithms rely heavily on the choice of starting population, and as we showed in an application to our thymus data, they may group cells in the same cell cycle phase despite transcriptional correspondence to different biological populations. Additionally, when there are groups of many distinct cells in general, a trajectory cannot be inferred altogether and a quantitative method for cluster comparison is still needed.

In conclusion, we propose average overlap as a metric for direct quantification of marker gene similarity with broad applicability in diverse biological settings suited to explore and clarify the heterogeneity of cells in most challenging contexts that arise from differentiating cells or involve clonal populations, such as those driving cancer progression.

## METHODS

### The sc_average_overlap Python package

We have implemented the average overlap metric in Python and have published the ‘sc_average_overlap’ Python package, which is available at https://github.com/chrisvthai/sc_average_overlap, designed to work seamlessly with Scanpy, a commonly used Python library for scRNA-seq analyses. The package includes functions for computing pairwise AO scores between two clusters, performing hierarchical clustering of cell populations, and storing the resulting dendrogram in Scanpy’s native AnnData object. For hierarchical clustering, the AO values are subtracted from one to generate a distance metric, since AO ranges between 0 and 1, with higher overlap scores meaning a lower distance, and vice versa.

### Benchmarking average overlap’s performance

To benchmark AO when used as the metric for hierarchical clustering of cell populations in single cell RNA-seq data, we utilized the *Zhengmix8eq* dataset, generated for the purpose of benchmarking clustering performance in scRNA-seq analysis^20^. We used only five T-cell populations, which were CD8 cytotoxic cells and CD4 populations including helper, memory, naïve, and regulatory cells. We processed the data using Piccolo^26^ to perform feature selection and normalization, selecting 3,000 highly variable genes for downstream analysis. We then performed PCA and Leiden clustering with a resolution of 2.0 using Scanpy. We performed this process and all future downstream analysis 100 times in total, each with a different random seed given to the Leiden algorithm.

We generated marker genes for each resulting single-cell cluster by performing differential expression analysis using the Wilcoxon rank-sum test and comparing each cluster’s gene expression versus the rest of the cells, ranked by significance. We then grouped single-cell clusters using hierarchical clustering. The metrics used for the hierarchical clustering included Pearson correlation, Spearman correlation, Kendall-Tau coefficient, and AO, and were calculated with either the set of all cluster marker genes, composed of the union of each unsupervised cluster’s top 5, 10, 25, 50, or 100 differentially expressed marker genes, or the values of the first 50 principal components. When using the set of all cluster marker genes, Pearson, Spearman, and Kendall-Tau coefficients were calculated using each cluster’s expression values for each gene, while AO was calculated using the rankings of the cluster marker genes. When using principal component values, only Pearson, Spearman, and Kendall-Tau coefficients were calculated.

Once hierarchical clustering was computed, the clusters were iteratively merged in an automated manner, guided by the pair of clusters with the highest AO similarity, until five final clusters remained, corresponding to the five represented T-cell populations in the *Zhengmix8eq* dataset. Performance was evaluated in two ways. Firstly, using the ground truth labels, we calculated adjusted Rand index (ARI), adjusted mutual information (AMI), and the Fowlkes-Mallows score (FMS). Secondly, based on labeling each of the five final clusters according to its most enriched-for ground-truth label, we evaluated the separation of CD8 populations from CD4 populations. For the metrics and feature-sets used, we counted the occurrences where no clusters enriched with CD8 cytotoxic cells were identified. Additionally, in each clustering iteration, we calculated the ratio of the mean pairwise distances between the CD8 population (if identified) and each CD4 population versus the mean pairwise distances among the CD4 populations. In each occurrence where no CD8 cluster was detected, we set the mean distance ratio between CD8 and CD4 groups to 1.0 to indicate no detected difference in separation between CD8 and CD4 groups.

### Thymocyte analysis using average overlap

We applied the AO metric to a previously published dataset of mouse thymocytes^28^. We first performed feature selection and normalization using Piccolo, excluding cells expressing more than 12% mitochondrial genes or 80% ribosomal genes and selecting 3,000 highly variable genes for downstream analysis. We performed Principal Component Analysis (PCA) and Leiden clustering at resolution 1.0 using Scanpy, and visualized the single cell clusters using Uniform Manifold Approximation and Projection (UMAP). We performed differential gene expression analysis using the Wilcoxon rank-sum test and assigned the top 25 differentially expressed genes, ranked by significance, as a set of marker genes in each cluster. As we were interested in T-cell populations only, we excluded small populations of granulocytes and B-cells detected through scoring against known markers of each cell type. We combined all clusters’ marker genes into a single set for calculating AO for subsequent calculation of pair-wise distances between the clusters and hierarchical clustering.

We obtained and visualized the expression of each set of cluster marker genes in 12 sorted populations of thymic cells in mice from the ImmGen project using the MyGeneSet tool^31^. To summarize the expression of each single-cell cluster’s marker genes in ImmGen sorted bulk populations, we first transformed the expression counts into a normalized z-score and then combined the transformed values into a single composite z-score using Stouffer’s method.

We used Scanpy’s score_genes function for scoring each cell according to phases of the cell cycle using a list of 97 known cell cycle marker genes^33^. We used the same scoring method to distinguish between DN3a and DN3b populations based on a list of genes upregulated in purified wildtype DN3a/b cells^35^.

## AUTHOR CONTRIBUTION

CT and HK conceived the study. CT performed the analytical experiments and conducted all analyses with contribution and input from AS. DH and HK supervised the work. All authors drafted the manuscript. All authors read and approved the final manuscript.

## SUPPLEMENTARY FIGURE LEGENDS

**Supplementary Figure 1. Distributions of pair-wise average overlap scores on randomly shuffled lists.** Across 2,000 iterations, randomly shuffling two ranked lists that contain the same set of elements yields average overlap distances that closely follow a normal distribution with a mean of 0.5 and a variance that is inversely correlated with the length of the list.

**Supplementary Figure 2. a**, Adjusted Mutual Information (AMI) and **b**, Fowlkes-Mallows index (FMI) performance measures based on the enrichment of ground truth labels in five derived cell populations, across different metrics and number of marker genes used in the T-cells in the *Zhengmix8eq* dataset.

**Supplementary Figure 3. Upregulation of DN3a- and DN3b-related genes in developing mouse thymocytes.** Cells were scored by expression of genes upregulated in purified wildtype DN3a and 3b cells, determined by study from Vogel et al^35^.

**Supplementary Figure 4. a**, The expression of top 5 marker genes for each annotated cell population. **b**, The expression of marker genes expressed in more than 50% of their respective cell population.

**Supplementary Figure 5. Pair-wise differential expression between apoptotic and normal DN4/ISP cells in G2/M phase**. Differentially expressed genes between these groups were determined with the Wilcoxon rank-sum test. Normalized expression of the top 20 genes upregulated in the apoptotic group are shown.

**Supplementary Figure 6. Diffusion pseudotime inferred on developing thymocytes**. Among the group of cells that map to double negative thymocytes, when specifying cluster 0 as the starting point, the inferred trajectory closely matches our annotated stages of development. However, when setting cluster 2 as the start, two additional groups of cells, clusters 7 and 8, also share the same pseudotime value of 0.

**Supplementary Figure 7. Hierarchical clustering of Leiden clusters in thymus data using Pearson correlation**. Pearson correlation is calculated with the same set of top 25 marker genes in each Leiden cluster as previous analysis using the average overlap metric. Grouping based on either cell cycle phase or neighboring stages of T-cell development is inconsistent.

