## Supplementary Figures for "Guiding clustering and annotation in single-cell RNA sequencing using the average overlap metric"

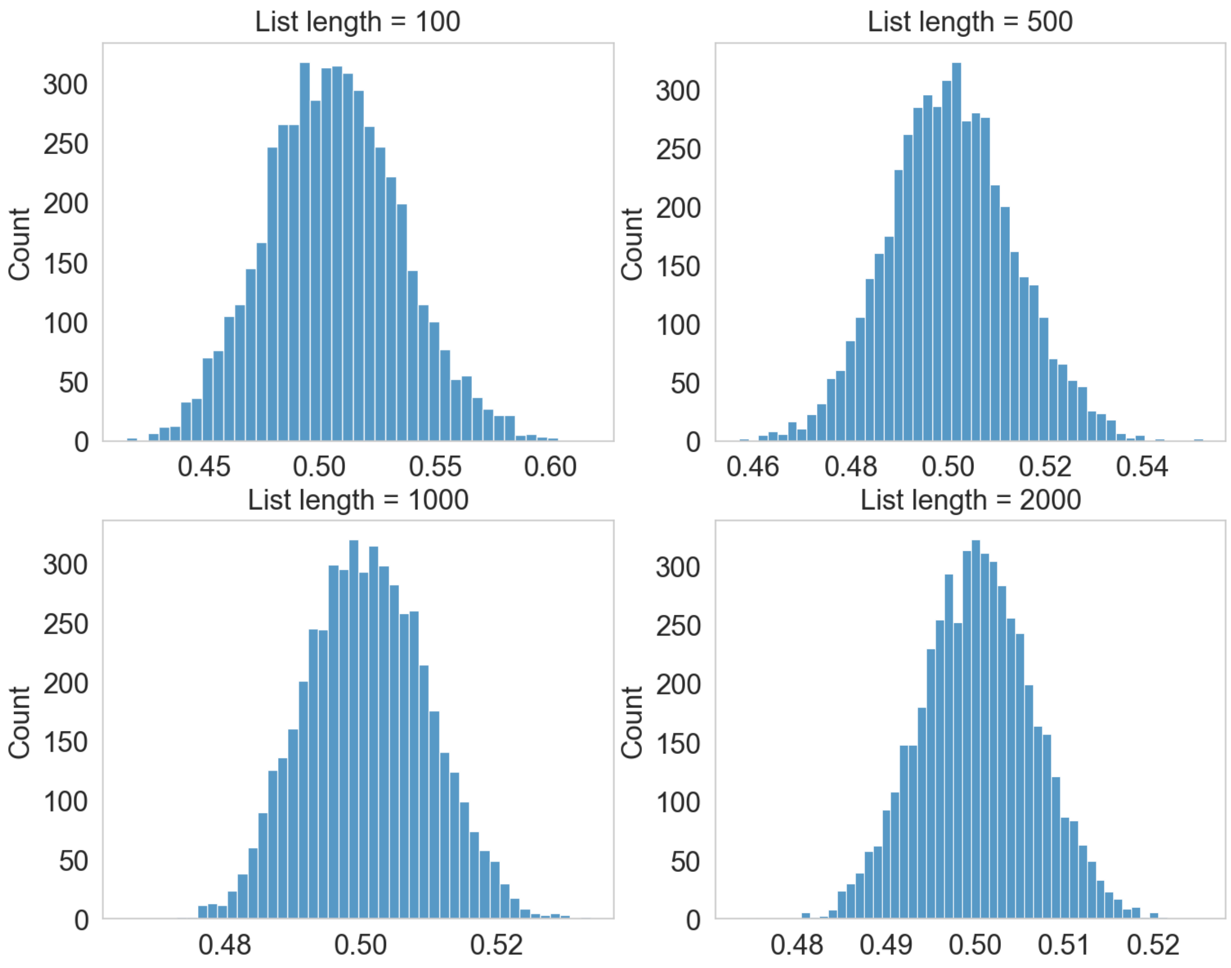


**Supplementary Figure 1. Distributions of pair-wise average overlap scores on randomly shuffled lists.** Across 2,000 iterations, randomly shuffling two ranked lists that contain the same set of elements yields average overlap distances that closely follow a normal distribution with a mean of 0.5 and a variance that is inversely correlated with the length of the list.

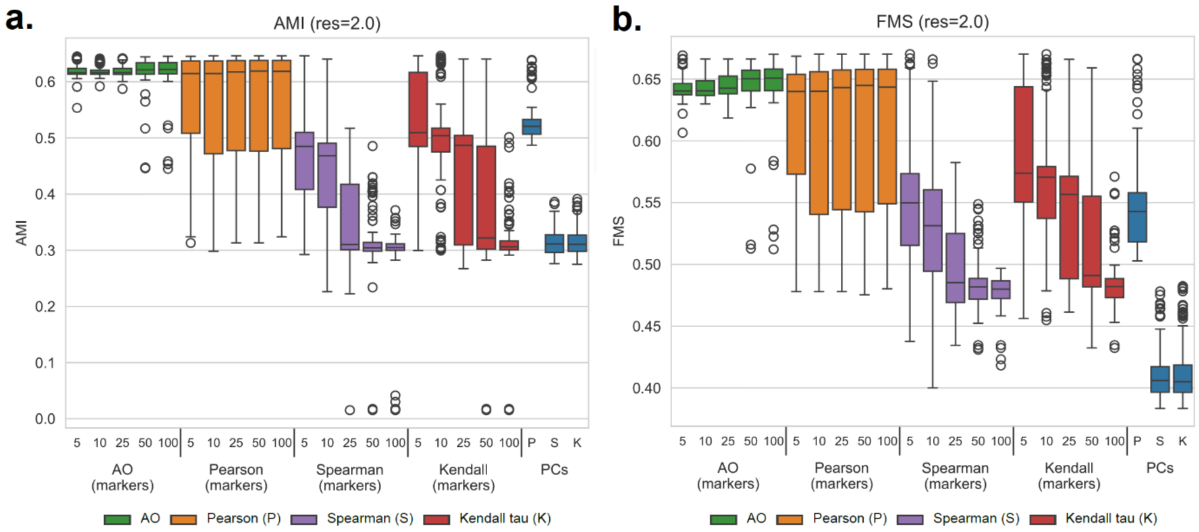


**Supplementary Figure 2. a,** Adjusted Mutual Information (AMI) and **b,** Fowlkes-Mallows index (FMI) performance measures based on the enrichment of ground truth labels in five derived cell populations, across different metrics and number of marker genes used in the T-cells in the Zhengmix8eq dataset.

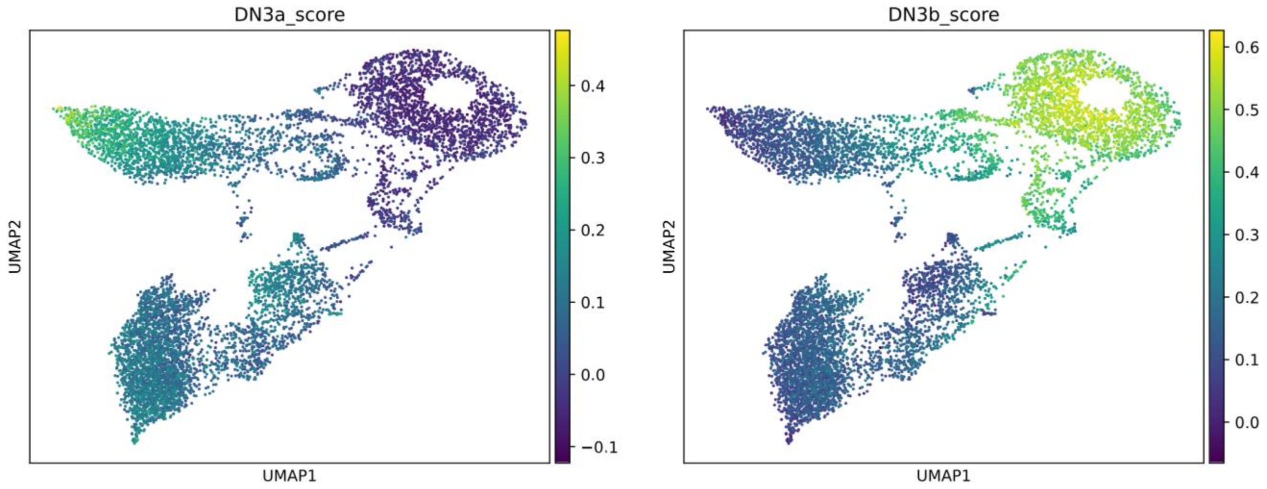


**Supplementary Figure 3**. **Upregulation of DN3a- and DN3b-related genes in developing mice thymocytes.** Cells were scored by expression of genes upregulated in purified wildtype DN3a and 3b cells, determined by study from Vogel et al[35].

**a**

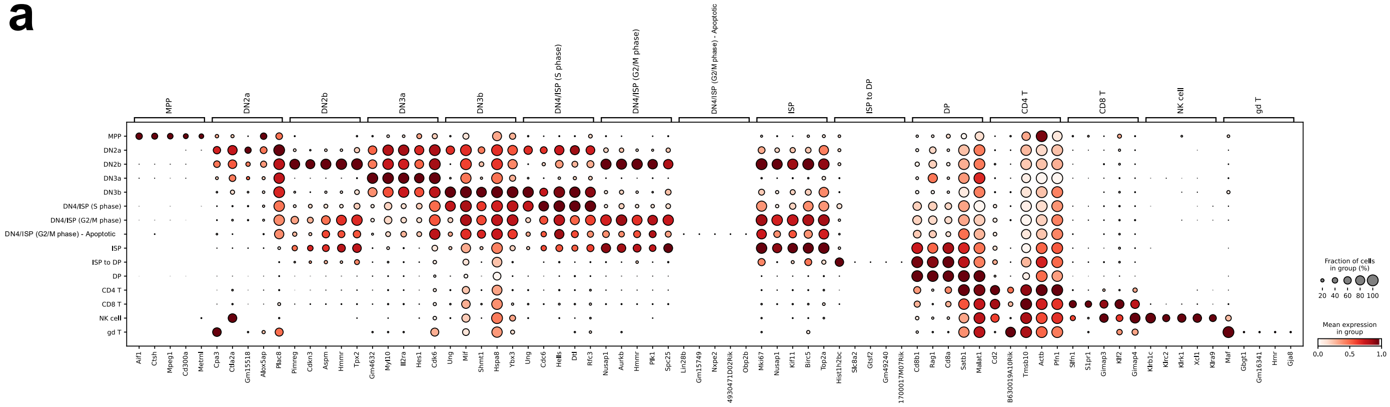


**b**

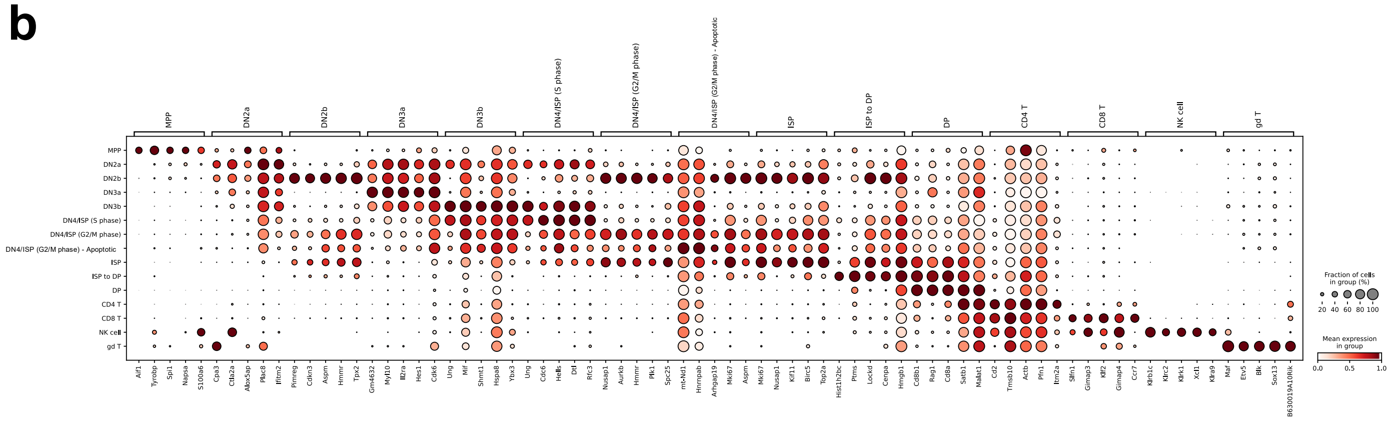


**Supplementary Figure 4. a,** The expression of top 5 marker genes for each annotated cell population. **b,** The expression of marker genes expressed in more than 50% of their respective cell population.

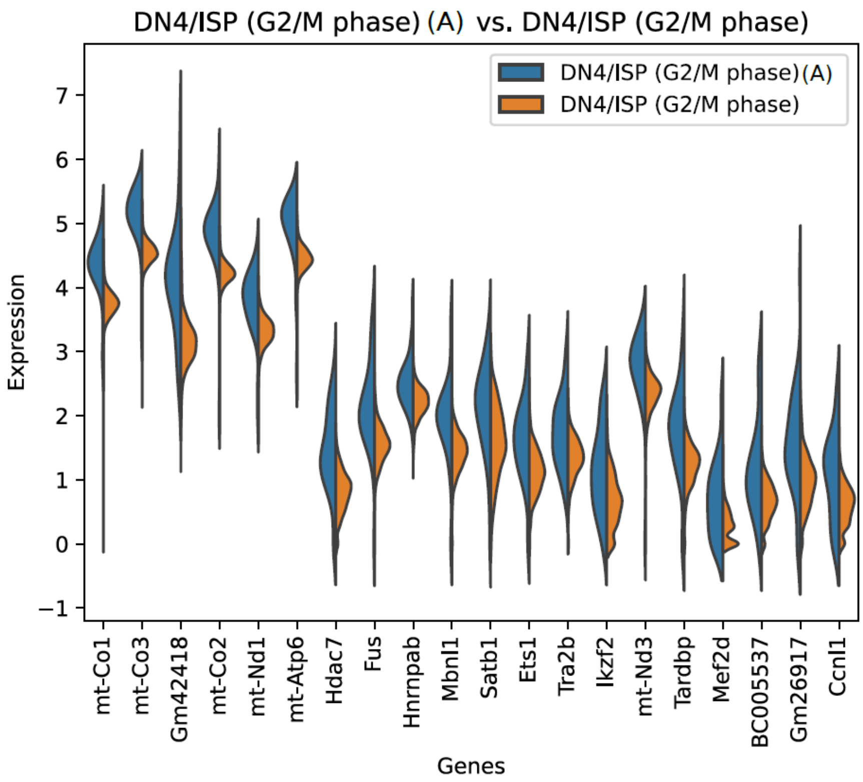


**Supplementary Figure 5. Pair-wise differential expression between apoptotic and normal DN4/ISP cells in G2/M phase**. Differentially expressed genes between these groups were determined with the Wilcoxon rank-sum test. Normalized expression of the top 20 genes upregulated in the apoptotic group are shown.

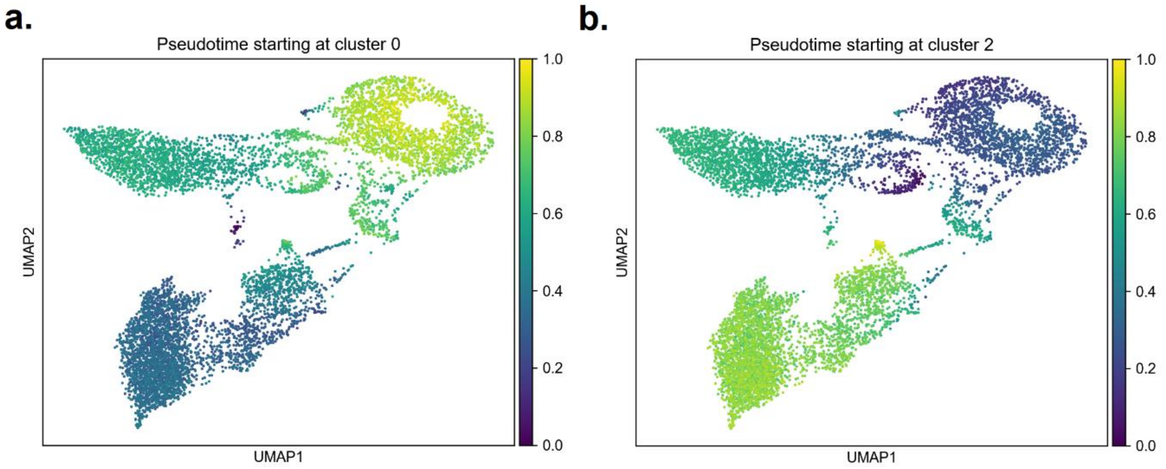


**Supplementary Figure 6. Diffusion pseudotime inferred on developing thymocytes**. Among the group of cells that map to double negative thymocytes, when specifying cluster 0 as the starting point, the inferred trajectory closely matches our annotated stages of development. However, when setting cluster 2 as the start, two additional groups of cells, clusters 7 and 8, also share the same pseudotime value of 0.

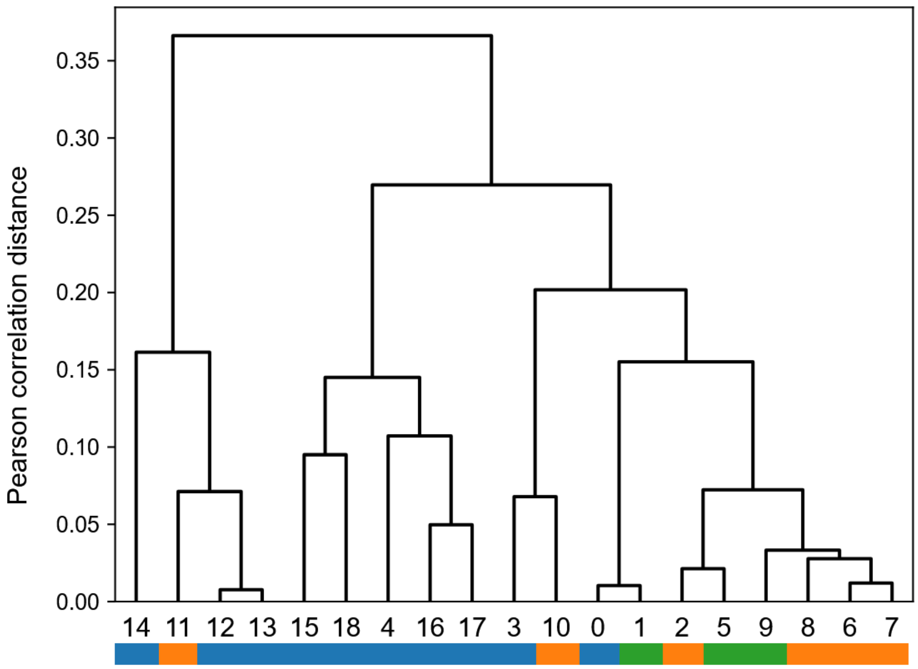


**Supplementary Figure 7. Hierarchical clustering of Leiden clusters in thymus data using Pearson correlation**. Pearson correlation is calculated with the same set of top 25 marker genes in each Leiden cluster as previous analysis using the average overlap metric. Grouping based on either cell cycle phase or neighboring stages of T-cell development is inconsistent.
